# The iPhylo Suite: An Interactive Platform for Building and Annotating Biological and Chemical Taxonomic Trees

**DOI:** 10.1101/2024.03.25.586513

**Authors:** Yueer Li, Chen Peng, Fei Chi, Zinuo Huang, Mengyi Yuan, Chao Jiang

**Affiliations:** MOE Key Laboratory of Biosystems Homeostasis & Protection, and Zhejiang Provincial Key Laboratory of Cancer Molecular Cell Biology, Life Sciences Institute, Zhejiang University, Hangzhou, Zhejiang 310030, China; State Key Laboratory for Diagnosis and Treatment of Infectious Diseases, First Affiliated Hospital, Zhejiang University School of Medicine, Hangzhou, Zhejiang 310009, China; Innovation Center of Yangtze River Delta, Zhejiang University, Hangzhou, Zhejiang, China; Center for Life Sciences, Shaoxing Institute, Zhejiang University, Shaoxing 321000, China

**Keywords:** Taxonomic analysis, Chemical taxonomy, Tree visualization, Interactive annotation

## Abstract

Accurate and rapid taxonomic classifications are essential for systematically exploring organisms and metabolites in diverse environments. Many tools have been developed for biological taxonomic trees, but limitations apply, and a streamlined method for constructing chemical taxonomic trees is lacking. We present the iPhylo suite (https://www.iphylo.net/), a comprehensive, automated, and interactive platform for biological and chemical taxonomic analysis. The iPhylo suite includes the Tree and Visual web-based modules for interactive building and annotating biological and chemical taxonomic trees, and a stand- alone command-line interface (CLI) module for local use and deployment on high-performance computing clusters. Furthermore, the iPhylo visualization module is fully implemented in R, allowing users to save progress locally and modify the underlying R codes. Finally, the iPhylo CLI module extends the analysis to all hierarchical relational databases. In summary, the iPhylo suite provides an integrated interactive framework for in-depth taxonomic analyses of biological and chemical features and beyond.

**Highlights:**

- A platform for both biological and chemical classification analysis and visualizations.
- A stand-alone Command line interface module for local or HPC deployment.
- R-based interactive visualization module with session-saving options.
- Supports customized hierarchical databases for analysis beyond taxonomy.

## Introduction

The systematic investigation of organisms and metabolites in human and diverse environments requires accurate and fast taxonomic classifications. Over the years, the comprehensive biological taxonomy, or Linnaean taxonomy, has been curated by NCBI [1] and EMBL [2] to organize the ever-increasing number of species. The fast-growing need for integrating metabolomic perspective in microbiome and precision medicine research provokes the need to analyze chemical taxonomy conveniently. Similar to the biological classification system, several chemical classification systems have been developed, including the ChEBI ontology [3], LIPID MAPS [4], and ChemOnt [5]. These systems use various hierarchical chemical criteria to organize chemical compounds. Utilizing a taxonomic tree structure to present a classification system is highly intuitive, allowing for a concise representation of the hierarchical relationships between features.

Several online and stand-alone tools offer taxonomic classification, visualization, and annotations in various combinations, including MEGA [6], ETE Toolkit [7], phangorn [8], taxtree [9], and PhyloT [10] for biological trees. The NCBI Taxonomy website offers an online tool (https://www.ncbi.nlm.nih.gov/Taxonomy/Browser/wwwtax.cgi) for constructing taxonomic trees. Qemistree [11], CluMSID [12], and BioDendro [13] are computational tools designed for clustering and identifying chemicals based on mass-spectrometry features, and they also provide the corresponding dendrogram visualization. ITOL [14] is a well-established web-based option for visualization and annotation tools, and the newly developed web-based tool TVBOT [15] offers similar functionalities.

However, several limitations apply. Software packages such as ETE Toolkit and phangorn require programming proficiency and have a steep learning curve for non-experts, and taxtree suffers from undesirable waiting time. The TaxBrowser of the NCBI website suffers from limited functions, unattractive interfaces, and incompatibility with batch submissions. The monetary requirement to use a few mainstream and user-friendly online tools is a severe limitation for researchers in less developed countries. In summary, most software implementations only apply to different stages of the entire workflow of taxonomic analysis. Switching between software may lead to data format incompatibilities and other issues. More importantly, while constructing phylogenetic trees is popular and relatively easy for biological features, constructing and annotating chemical taxonomic trees is difficult due to a lack of a convenient method. With the growing need for integrating metagenomic and metabolomic data in precision medicine and microbiome research, a unified platform for building and annotating biological and chemical taxonomic trees is imperative. Most interactive tools do not have the free option for saving and uploading tree-building sessions for more complex projects. Finally, no current platform offers the underlying visualization codes in R, a popular language for data analysis. For advanced users, accessing the R source codes for plotting and annotation is highly desirable for further customizations.

We present the iPhylo suite: a fully automated and interactive platform for biological and chemical taxonomic analysis. The iPhylo suite includes three modules. Two web-based modules, iPhylo Tree and iPhylo Visual, aim to streamline the workflow, encompassing tree generation, interactive tree visualization, and extensive graphic and textual annotations. A stand-alone module, iPhylo CLI, was designed for local use and high-performance computing applications and equipped with up-to-date biological and chemical taxonomic databases.

## Methods

### The overall workflow of the iPhylo suite

The iPhylo suite features three modules (Figure 1): (1) The iPhylo Tree rapidly generates biological or chemical taxonomic trees for up to tens of thousands of organisms and chemicals within minutes (https://www.iphylo.net/). (2) The iPhylo Visual was developed based on the R framework [16–18] for visualizing and extensively annotating taxonomic trees (https://www.iphylo.net/visual/). The iPhylo Visual also offers the convenience of saving and uploading work sessions locally, as well as access to source codes for plotting and annotating. Importantly, the Tree and Visual modules are seamlessly integrated, enabling users to swiftly import trees and leaf data from construction to annotation with a single click. (3) The iPhylo CLI is an offline command-line version of the Tree module with integrated databases that can be deployed locally or on high-performance computing clusters. Importantly, the iPhylo CLI can also construct customized taxonomic trees based on a user-defined hierarchical database so the applications of the iPhylo suite can be extended beyond biological and chemical classifications.

**Figure 1.**
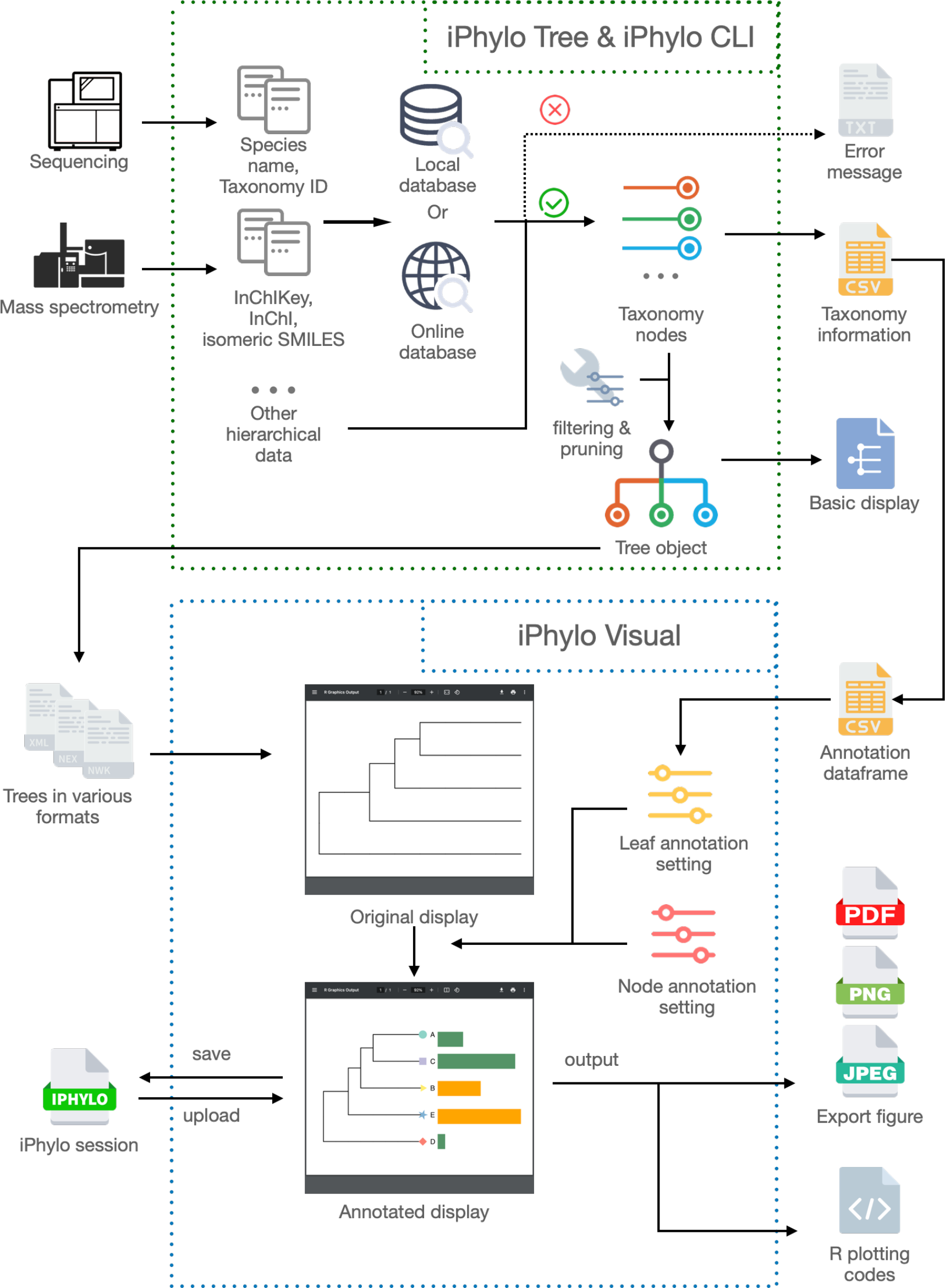
The overview of the iPhylo suite The iPhylo Tree, CLI, Visual modules, and the entire workflow, accompanied by the different formats of input and output files.

### The Intuitive Interface

The iPhylo suite server employs user-friendly interfaces, including but not limited to the navigation home, the tree construction page, the annotation dashboard, and the inspection pages (Figure 2). The homepage displays an array of visually captivating dynamic effects that introduce the website’s unique features and functionalities (Figure 2A). The tree-constructing (Figure 2B) and annotation dashboard (Figure 2C) pages feature intuitive and functional form layouts. The inspection page adopts the dragging selector to convert the branches and nodes into structured data, facilitating the exploration of any part of the tree (Figure 2D). There are dedicated tutorial help pages and a gallery page showcasing examples.

**Figure 2.**
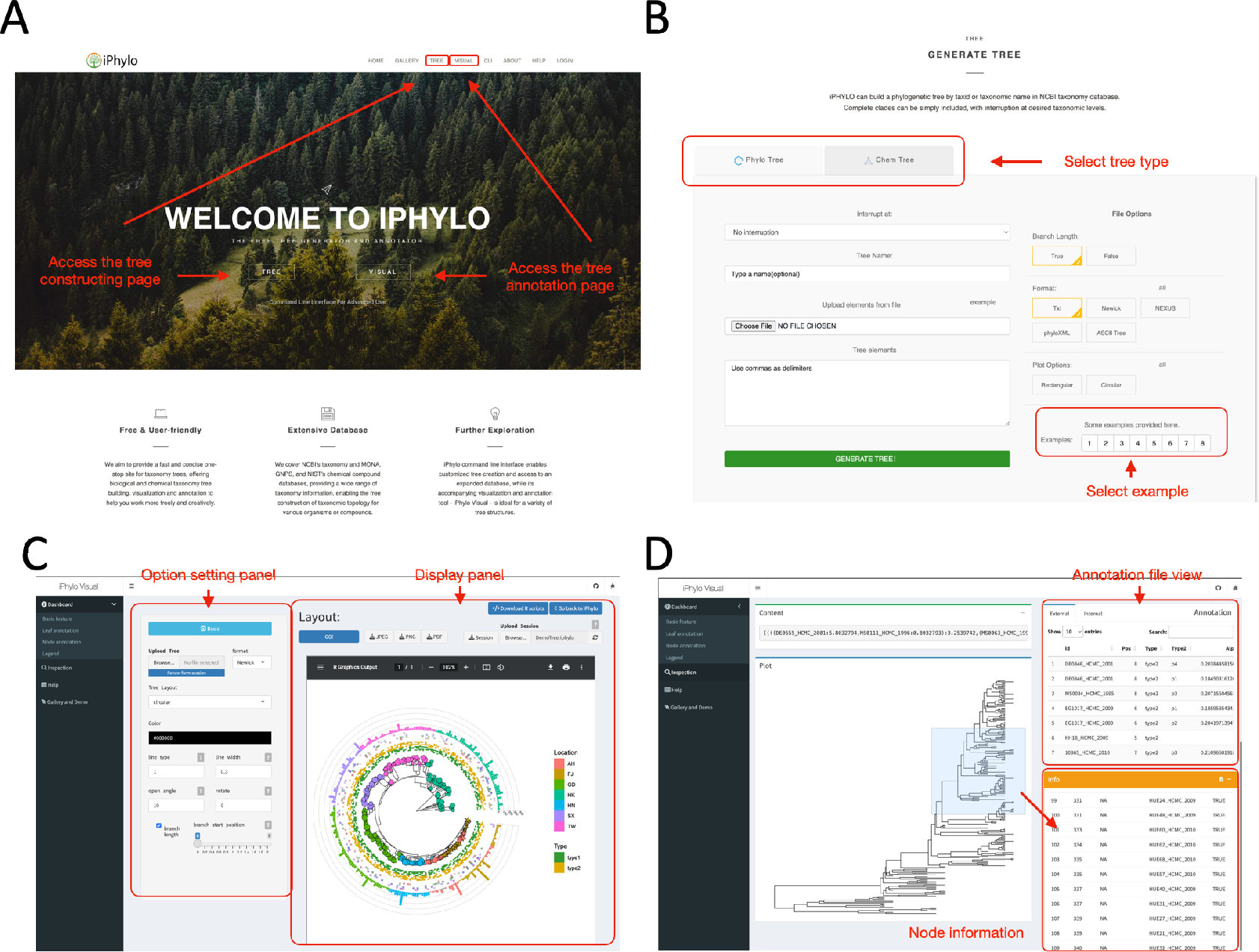
The user interface of the iPhylo suite **A**. The home page. **B**. The tree construction page. **C**. The annotation dashboard page. **D**. The annotation inspection page.

### iPhylo Tree

The iPhylo Tree module is an online web application for constructing taxonomic trees from biological species or chemical compounds. Historically, constructing phylogenetic trees may involve features such as morphology [19,20], biochemical [21], or behavioral features [20]. Modern phylogenetic trees are inferred from DNA or protein sequence alignments using tools such as BEAST [22] and MEGA [6]. However, these alignment-based tools cannot easily construct trees spanning all life domains, require heavy data input, and can be time-consuming. Additionally, The results of chemical trees using methods based on molecular similarity, such as Qemistree [11], CluMSID [12], and BioDendro [13], start from spectrum data and depend on the clustering algorithm. Therefore, iPhylo Tree presents a swift and efficient alternative. It builds biological taxonomic trees based on the NCBI Taxonomy, enabling rapid retrieval of lineage information spanning all life domains. Similarly, it creates chemical taxonomic trees by utilizing ChemOnt, thereby eliminating the need for extensive input and the impact of the algorithm. A complete pipeline of the iPhylo Tree includes the following steps:

*Step 1:* For constructing biological taxonomic trees, users provide a set of species names or corresponding taxonomy identifiers (TaxID). The species names are relatively flexible as long as they are indexed in the NCBI taxonomic database, and “scientific name,” “synonym,” or “common name” are all acceptable. For chemicals, the input can be InChIKey, InChI, and isomeric SMILES. Currently, common chemical names are not supported due to extensive heterogeneity and unnamed chemicals. The InChIKey is highly recommended because some chemicals lack InChI and isomeric SMILES information.

*Step 2:* Taxonomy querying involves querying input biological and chemical features against the iPhylo databases to extract taxonomic data. The iPhylo databases comprise biological and chemical divisions. The biological database includes the entire species classification information sourced from the NCBI taxonomy database, encompassing a total of 2,388,300 TaxIDs. The chemical database includes classification details for 801,308 functional compounds compiled from MONA, GNPS, and NIST databases, with a focus on human metabolome. The chemical taxonomic data is managed based on the structure- based chemical classification methodology called ChemOnt [5]. Moreover, the databases will be regularly updated to keep up with the expanding knowledge of biological and chemical features.

*Step 3:* To construct taxonomic trees, we used an object-oriented method to transform queried taxonomic data into an n-ary tree object, enabling easy representation in formats such as Newick. For output, we employed a depth- first post-order traversal algorithm to navigate the tree, preserving its topology as a Newick format string. Subsequently, iPhylo utilizes the phylo package from BioPython [23] to transform the Newick-formatted string to other formats, including Nexus and PhyloXML. Additionally, we provide an ASCII representation and a PDF visualization of the tree generated by the ggtree [17] package.

The resulting biological taxonomic tree is organized by a unified number of taxonomic levels: Domain, Kingdom, Phylum, Class, Order, Family, Genus, and Species, excluding informal ranks like subphylum and subclass. The chemical taxonomic tree is also organized in a similar structure, from the highest to the lowest: Kingdom, SuperClass, Class, SubClass, Parent Level 1, and Parent Level 2.

Furthermore, iPhylo Tree is capable of filtering the tree by interrupting each branch at a specified level. The “|subtree” operator enables the extraction of a group of taxa that includes the common ancestor and all descendants within the specified clade, such as “Primates|subtree” or “Hydroxyindoles|subtree”.

### iPhylo CLI

The iPhylo CLI serves as an extension of the web-based iPhylo Tree. The iPhylo CLI includes four modules, namely (1) the phylo tree module, (2) the chemical tree module, (3) the chemical online module, and (4) the csv2tree module. The phylo tree module and chemical tree module offer the same functionality as the online iPhylo Tree but run locally. Notably, the exclusive online chemical module enables chemical information retrieval from ClassyFire [5] API, including the chemicals processed with corresponding taxonomy data stored in ClassyFire’s database. Over 70 million chemicals can be queried through this online module, which is continually growing. This feature significantly augments the capabilities of chemical taxonomy analysis. Additionally, the csv2tree module empowers users to create customized trees from CSV files directly.

The iPhylo CLI is tailored for efficiency and scalability. After its initial execution, which downloads the necessary database resources, the subsequent runs operate in an offline local mode. This design ensures rapid execution and is well-suited for deployment on high-performance computing clusters (Supplementary Figure S1).

### iPhylo Visual

The iPhylo Visual is an interactive online tool designed to facilitate the display, annotation, and inspection of tree-based structures, including but not limited to phylogenetic and chemical taxonomic trees generated from iPhylo modules.

The iPhylo Visual simplifies the process of annotating taxonomic trees by adopting a data frame-compatible format, enabling users to encapsulate all required information within one data frame for leaf annotation and one data frame for node annotation, respectively. Within the data frame, rows correspond to tree nodes, and columns represent specific features. Users can efficiently navigate and manage these uploaded data frames through the provided online table viewer, with sorting and retrieval capabilities. This design avoids uploading multiple annotation files and is directly compatible with R.

The iPhylo Visual leverages the full graphical capabilities of ggtree [17] and ggtreeExtra [18] for visualizing, manipulating, and annotating tree-structured data. The iPhylo Visual provides the “Basic feature”, “Leaf annotation”, “Node annotation”, and “Legend” tabs for annotation controls. Users can choose from multiple layouts (e.g., circular, rectangular; Supplementary Figure S2), with adapting annotations. Customization options include branch thickness, color, and angle. The “Leaf Annotation” and “Node Annotation” tabs provide methods for adding specific markers to leaves and nodes, respectively (Supplementary Figure S3). Additionally, the “Legend” tab allows users to adjust the canvas size and legends for each annotation layer.

The iPhylo Visual emphasizes the ease of exporting and replicating tree displays with the “.iphylo” session files, packing all essential data for generating trees and maintaining compact sizes, e.g., 30 kB for a 1000-species tree. All sessions can be saved to and uploaded from the local computer, enabling easy storage, management, and collaboration.

iPhylo Visual supports exporting trees in PNG, PDF, and JPEG formats. Its “Export Code” feature allows a one-click download of visualization data, including the tree, annotations, parameters (JSON), and an R script. Running this script locally replicates the visualization, and users proficient with R programming can further modify the script.

### Implementation

The iPhylo Tree web server stores taxonomy data with MariaDB 5.5.60 database. The back-end of the iPhylo Tree webserver was developed using Python and the Flask framework version 2.2.2. The iPhylo Visual uses the R/Shiny framework for advanced tree display and annotation. We employed the Twitter Bootstrap template version 4.6 for the front end to create a visually appealing and user-friendly graphical interface. The web-based applications are deployed on a server with 32-core CPUs (2 GHz), 96GB RAM, and 1TB of storage to ensure that the hosting is scalable with the large increase in use. The iPhylo CLI was developed alongside the server version, with Python 3.8.15 and SQLite 3.39.3.

### Results management

The iPhylo suite offers free usage, including for commercial purposes, and does not require user registration. Besides, the iPhylo Tree module provides users with a login option to automatically save their generated tree history records. Notably, the registration does not require sensitive personal information, such as email addresses. The data for visualization and annotation in iPhylo Visual is managed with sessions. With the initiation of each new session within the user’s web browser, we establish temporary storage on our server, each associated with a unique identifier, enabling user download activity. When the session terminates, this data is cleared to ensure privacy and security.

## Results

To demonstrate the applications of the iPhylo suite, we constructed, visualized, and annotated a few phylogenetic and chemical taxonomic trees with data from published studies. The source data and iPhylo Visual session files for all case studies are available on the iPhylo website.

### Case Study 1: Visualization of the genome catalog of diverse bacterial strains from a glacial microbiome dataset

Studying glacier microorganisms facilitates the exploration of glacier microbial diversity, functionality, and evolution while assessing potential environmental and health risks linked to glacier melting [24]. In this example, iPhylo was used to visualize the Tibetan glacier metagenome-assembly genomes (MAGs) [25] and cultivated bacterial genomes. The data consisted of 3,241 Tibetan glacier genomes, clustered into 968 species-level operational taxonomic units (OTUs). We utilized the iPhylo CLI to construct a customized tree that incorporated the GTDB classification for each genome provided in the original study. To optimize the visualization process without compromising the display and functionality of the system, we sampled and formed a subtree from the full tree. This optimization resulted in a phylogenetic tree with 440 tips and 174 internal nodes. We next used iPhylo Visual to display and annotate the tree with various genome characteristics (Figure 3A), including the source of the genome, presence of 16S rRNA, number of tRNA genes, genome size, GC proportion, and genome quality standards based on the MAG [26]. The resulting figure displayed the phylogenetic trees in an accurate and visually appealing manner, incorporating multiple feature annotation tracks.

**Figure 3.**
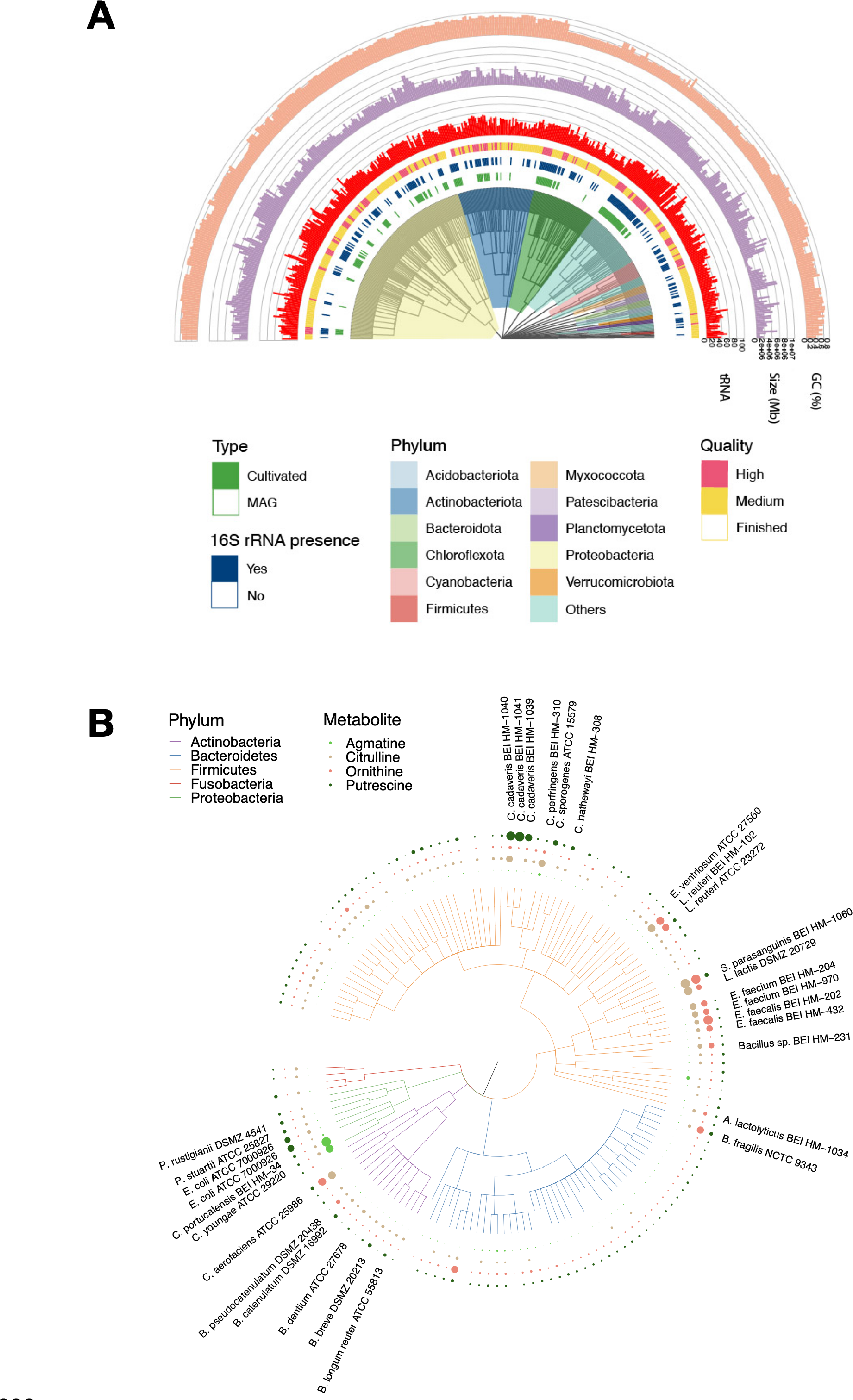
iPhylo visualizes phylogenetic trees with extensive annotation tracks **A**. A representative phylogenetic tree was constructed using the taxonomy data and visualized with annotations of various genome characteristics. Data source: Liu, Ji et al. 2022 [25]. **B**. A phylogenetic tree constructed using NCBI taxonomy IDs, with branches color-coded by phyla. The tree was further annotated with the abundance of four key metabolites. Data source: Han et al. 2021 [31].

### Case Study 2: Visualization of metabolic profiles for individual bacterial strains in the gut microbiome

Gut microbes are associated with multiple metabolic pathways and play a significant role in modifying host phenotypes and overall health [27,28], and can be affected by constant exposure to biological and chemical agents from diverse environmental sources [29,30]. Han et al. [31] reported a comprehensive metabolic profile of gut microbes encompassing 158 microbial strains and 833 metabolites. We selected 154 strains (taxonomy supported by NCBI Taxonomy) and the metabolites they produced as example data to illustrate the metabolite biosynthetic profiles of strains and the chemical taxonomy of the metabolites.

We first constructed a phylogenetic tree for the bacterial strains and annotated the phylum taxa. Next, we selected specific metabolites, namely putrescine, guanidine, ornithine, and citrulline, from the comprehensive pool of metabolites, as these four metabolites were explicitly noted in the origin study as potential indicators for measuring metabolic phenotypes of strains. We used iPhylo Visual to plot the abundance of these selected metabolites in all strains using bubble plots, as depicted in Figure 3B. Figure 4A illustrates the chemical taxonomic tree of the 813 metabolites successfully queried in the ClassyFire database, providing an ontology overview of the 11 SuperClasses, 84 Classes, and 157 SubClasses of chemicals.

**Figure 4.**
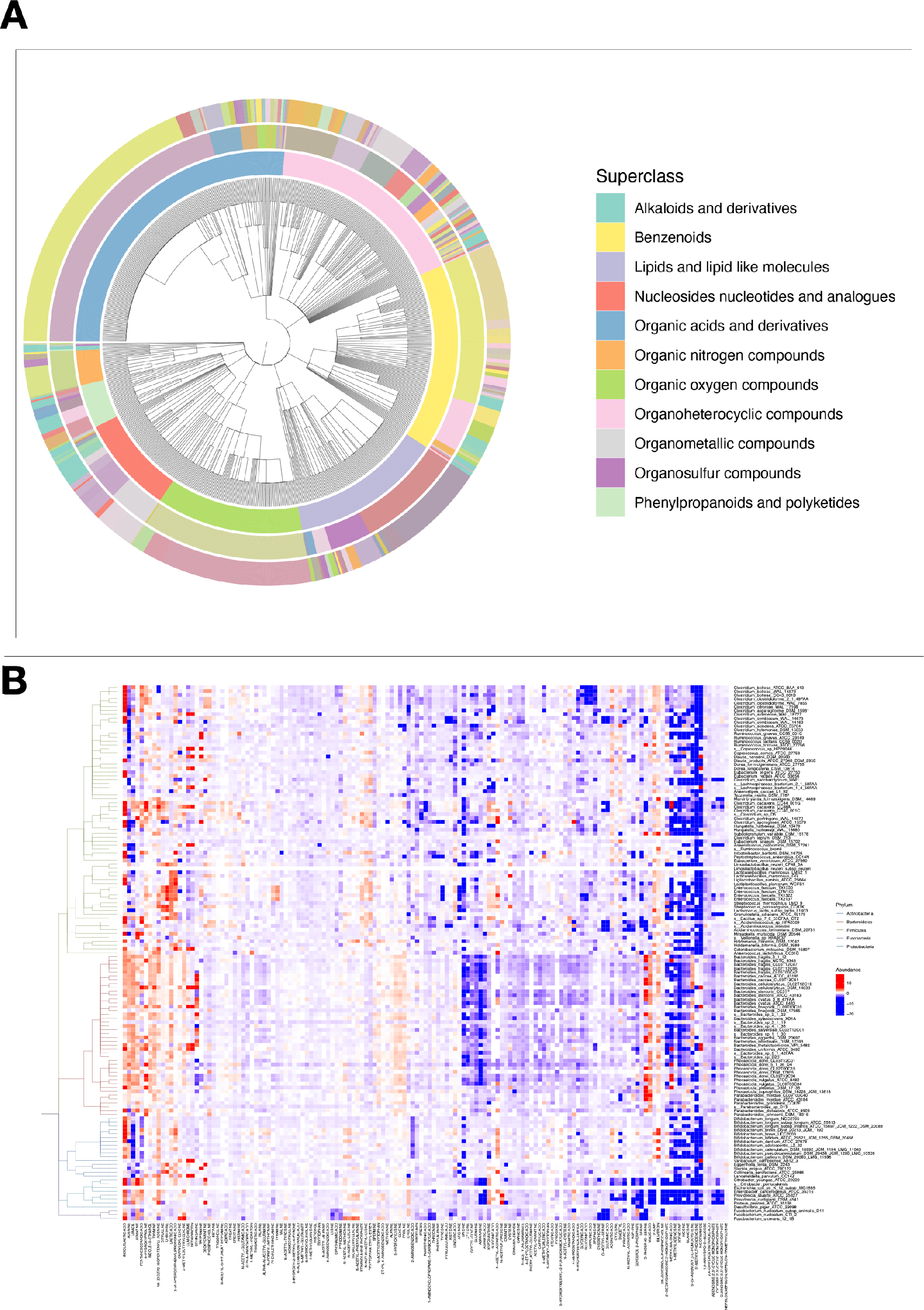
iPhylo visualizations of chemical database and heatmap **A**. A representative chemical taxonomy tree generated based on chemical InChIKeys and annotated at the superclass, class, and subclass levels (from inside to outside, only legends for superclasses are shown). Data source: Han et al., 2021 [31]. **B**. Heatmap visualization in iPhylo Visual: Phylogenetic tree of bacterial strains and the abundance of the associated metabolites. Colored branches represent different phyla. Data source: Han, S. et al. 2021[31]

### Case Study 3: The Versatility of iPhylo Visual

Heatmaps are important visualization methods in data analysis. In biological contexts, they depict associations between organisms and chemicals and may indicate the potential taxa interactions [32]. In this case study, we employ the heatmap to illustrate the taxonomy of microbes and their full metabolomic profiles, using the data in case study 2 (Figure 4B). The annotation file for generating a heat map requires the two-dimensional microbe-metabolite data table to be converted to a long-format table similar to the result of the “melt” function in R (as in the example table in Table S1). Due to the inherent nature of iPhylo Visual as a tool primarily designed for taxonomic trees, it only clusters the rows, or microbes in this case. Consequently, users should provide the clustering configurations for the columns, or in this case, metabolites.

This special case demonstrated the versatility of iPhylo Visual beyond tree visualization by incorporating heatmaps, box plots, violin plots, etc., as additional annotation layers to the trees.

### Case Study 4: A user-customized tree of statistical analysis methods

The iPhylo suite is compatible with any customized hierarchical relational database, as well as user-defined trees. Statistical methods encompass various classifications, such as hypothesis testing, ordination, factor analysis, clustering analysis, and correlation analysis. For demonstration purposes, we have selected several commonly used statistical methods based on their applicable data types and analysis goals. The hierarchical relationships for these methods were organized in CSV format (Supplementary Table 2) and processed by iPhylo CLI, resulting in a tree of common statistical methods. The tree was subsequently visualized in iPhylo Visual (**Figure 5**).

**Figure 5.**
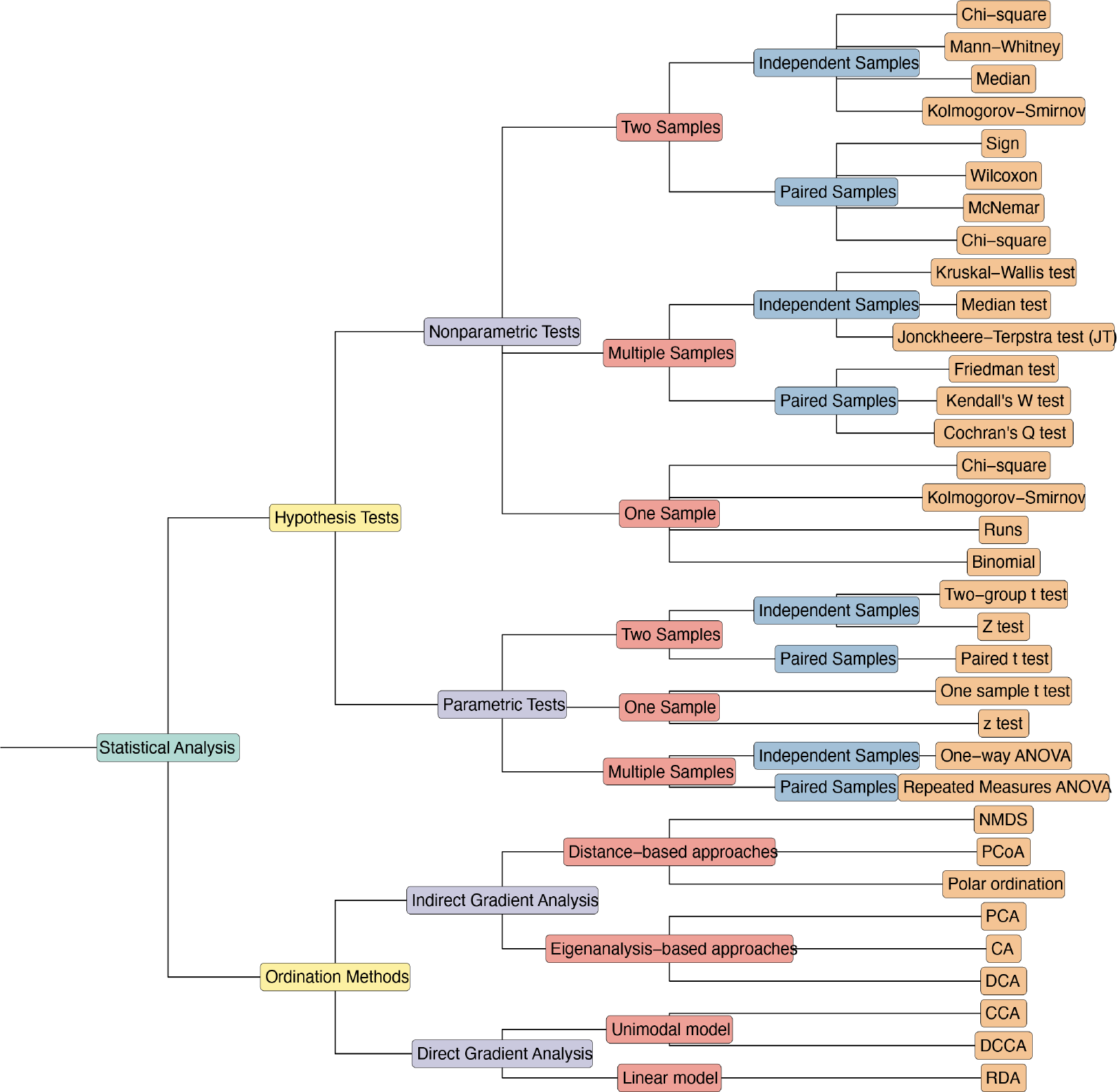
iPhylo visualizes user-customized trees A tree of common statistical methods was visualized as an example. We employed the iPhylo CLI module to organize the classifications of these statistical methods into a hierarchical tree structure and displayed it using iPhylo Visual. The tree is annotated with node labels that are color-coded based on taxonomic ranks.

## Discussion

Several existing tools for biological taxonomic analysis have been developed. However, no single tool or platform offers fully streamlined taxonomic analyses for biologicals and chemicals, and some tools are behind the paywall. The iPhylo suite aimed to address these issues by providing a fast and convenient solution. The iPhylo suite was developed with several principles: (1) Cross- platform compatibility. The iPhylo suite offers web-based and command-line services, which can be easily accessed across Windows, Mac OS, and Linux. (2) Integration. iPhylo seamlessly integrates the entire workflow of constructing, visualizing, and annotating taxonomy trees without switching platforms, reducing the room for error. (3) Customizability. iPhylo offers extensive customization options and extends its utility to all hierarchical relational databases. The data frame format was adopted for annotation to minimize the input requirements from users. Additionally, users can download and upload tree-building sessions and access the underlying R plotting code for further modifications.

Currently, the embedded chemical database focuses on functional compounds, representing only a small fraction of the entire universe of chemical substances. We provided a solution by querying online data resources in iPhylo CLI, but the advantage of high querying speed in the locally embedded chemical database cannot be replaced. Furthermore, other databases, such as the GTDB dataset, also present opportunities for future expansion. Looking forward, the iPhylo suite’s development roadmap includes plans to incorporate a broader range of databases not limited to biologicals and chemicals. We believe the iPhylo suite will greatly facilitate the general adoption of biological and chemical taxonomic analyses in diverse fields such as microbiome, metabolome, precision medicine, ecology, and environmental science.

## Data availability

All the demonstration data for the case study can be accessed on the iPhylo Visual website’s Help page (https://iphylo.net/visual). The iPhylo database is available for download as a “.db” file from the following URLs: Biological database: https://iphylo.net/resource/iphylo_db Chemical database: https://iphylo.net/resource/ichem_db

## Code availability

The source code for iPhylo CLI is open-source and can be found on GitHub (https://github.com/ARise-fox/iPhylo-CLI). Additionally, the raw data and corresponding preprocessing R script can be downloaded from the GitHub repository (https://github.com/ARise-fox/iPhylo-CaseStudyData).

## Authors’ contributions

YL and CJ conceived the study. YL developed the iPhylo Tree and CLI Module. YL and CP developed the iPhylo Visual module. FC provided the deployment on the computing cluster. ZH and MY collected the data for the chemical database. YL and CJ drafted and revised the manuscript with input from other authors.

## Competing interests

The authors have declared no competing interests.

## Acknowledgements

We are grateful to our colleagues at the core facility of the Life Sciences Institute, especially the NECHO high-performance computing cluster. This research was partly supported by the grant from NSFC (82173645).

## Supplementary material

Figure S1 Runtime results of generating tree in all formats Using a series number (100, 1000, 2000, 5000) of taxIDs or InChIKeys from the database as input data. Tests for each group were replicated 20 times.

Figure S2 iPhylo visualizations of user-customized trees (a) circular layout. (b) inward circular layout. (c) daylight layout. (d) rectangular layout. (e) slanted layout. (f) ellipse layout. (g) roundrect layout.

Figure S3 The leaf annotation types of iPhylo Visual (a) tile track. (b) bar track. (c) box track. (d) violin. (e) tip track. (f) point track. (g) tip label track.

Figure S4 The node annotation types of iPhylo Visual (a) strip track. (b) color branch track. (c) highlight track.

Table S1 Long-format table format for annotation in iPhylo Visual

Table S2 The hierarchical classification for common statistical methods in iPhylo CLI

