## Supplementary figures and images for "The iPhylo Suite: An Interactive Platform for Building and Annotating Biological and Chemical Taxonomic Trees"

### Supplementary Figure 1

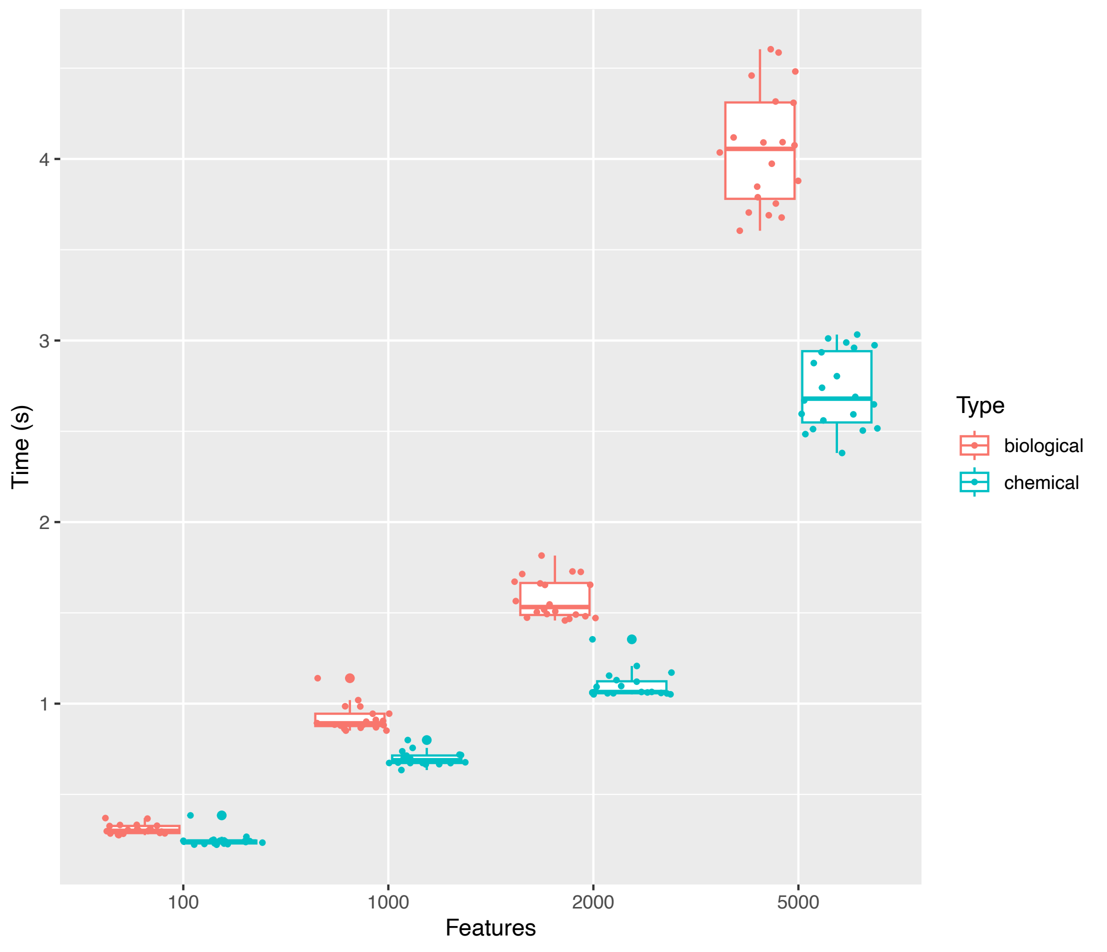

### Supplementary Figure 2

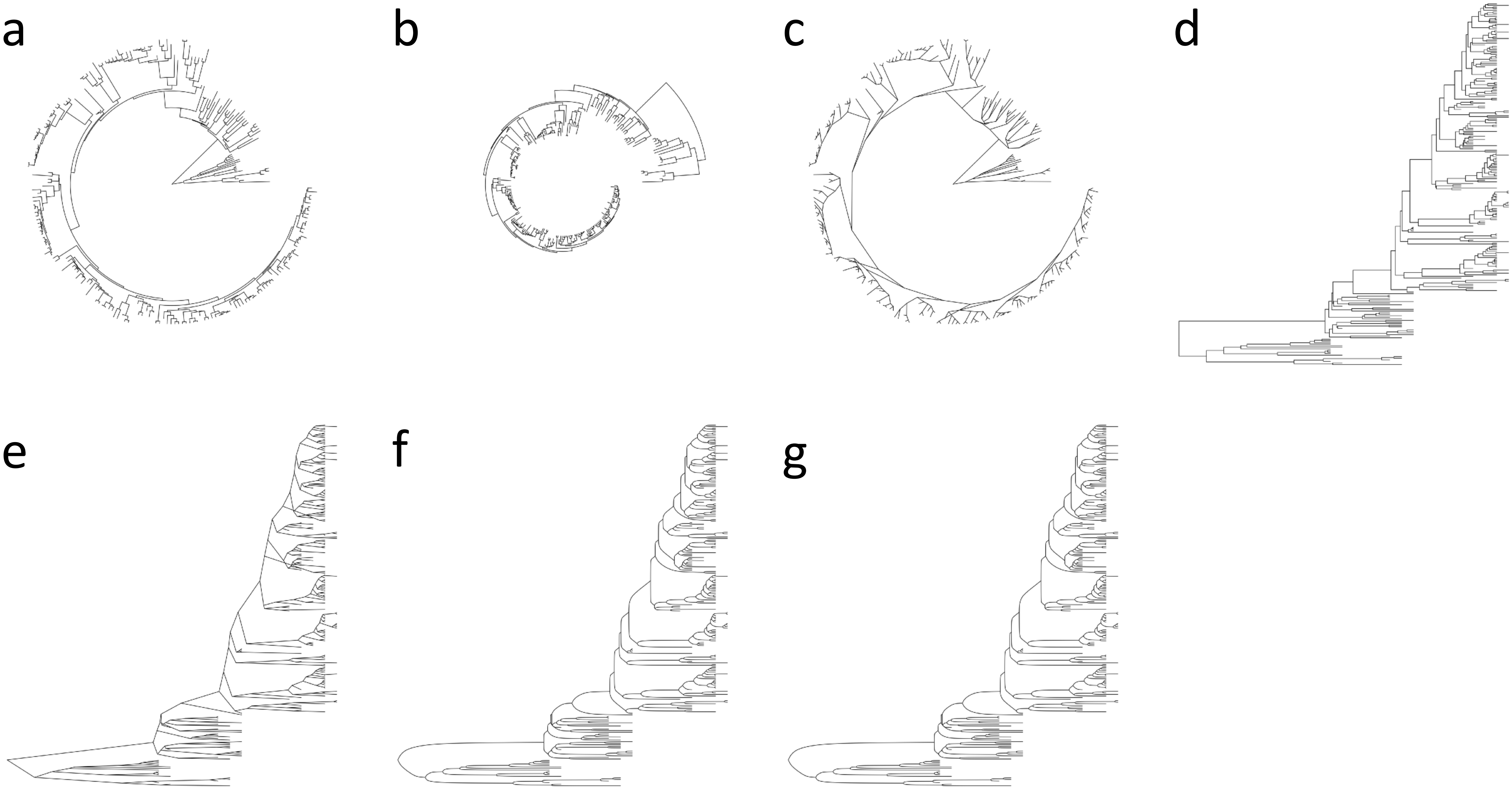

### Supplementary Figure 3

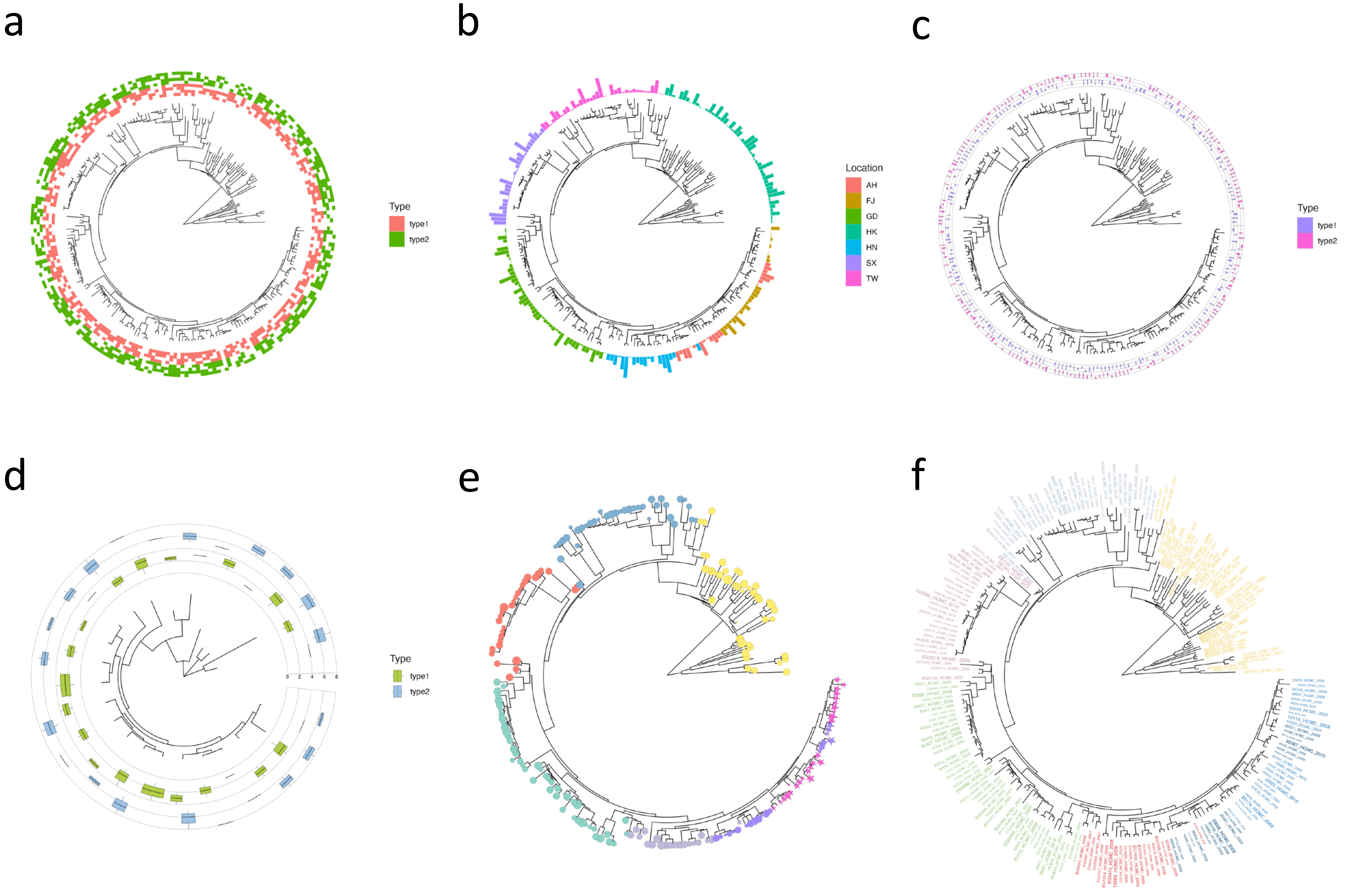

### Supplementary Figure 4

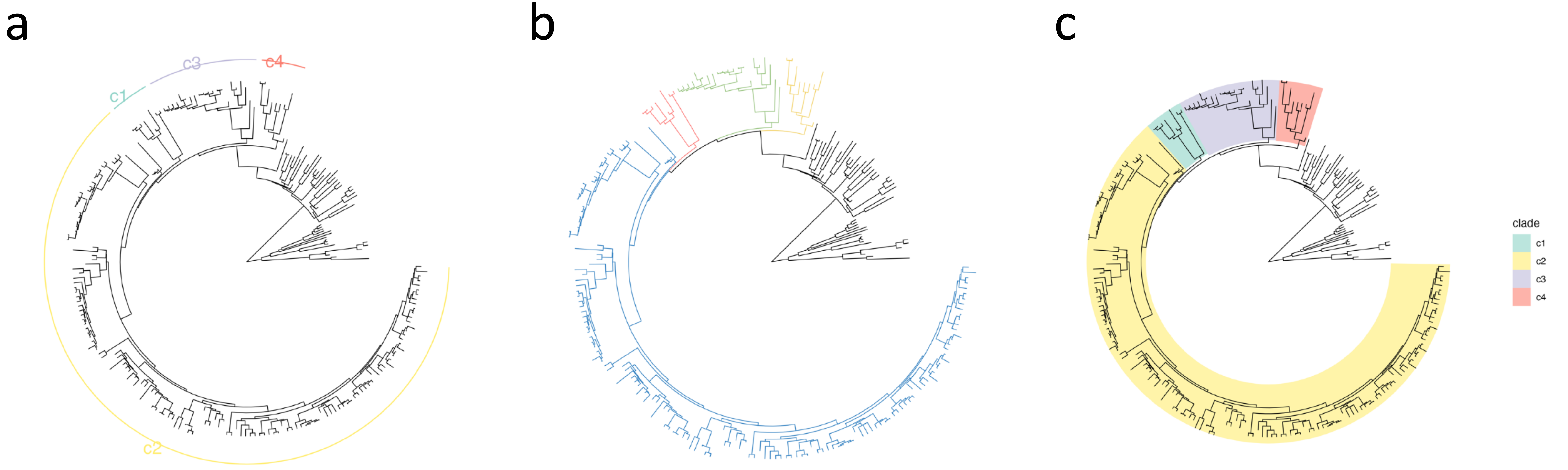
